# Changes in perceptual sampling contribute to representational drift

**DOI:** 10.64898/2026.06.24.734121

**Authors:** Yixin Yuan, John T. Serences, Mikio C. Aoi

**Affiliations:** Neurosciences Graduate Program, UC San Diego; Department of Neurobiology, UC San Diego; Halıcıoğlu Data Science Institute, UC San Diego; Department of Psychology, UC San Diego

**Author notes:** These authors contributed equally to the senior supervision of this work (John T. Serences, Mikio C. Aoi).

## Abstract

Gradual changes in neural response patterns to the same stimulus over time, termed *representational drift*, have been widely observed across cortical areas. Drift is typically attributed to intrinsic neural dynamics driven by synaptic plasticity or turnover. Here we test a complementary hypothesis that drift arises due to systematic changes in behavior such as shifts in attention or gaze. We conducted a longitudinal eye-tracking experiment in which fourteen adults freely viewed naturalistic images across 6 experimental sessions spanning 2–4 weeks, with a subset of images repeated within and across sessions. For each participant, we quantified the similarity of fixation density maps between session pairs using the Wasserstein distance. Fixation patterns became increasingly dissimilar with greater temporal separation between sessions, indicating systematic and directional drift in gaze behavior. To assess whether these behavioral changes could plausibly induce changes in neural representations, we passed fixation-masked images through CORnet-S, a hierarchical deep neural network model of the primate ventral visual stream. Representational distances, quantified as the Frobenius norm of pairwise activation differences, increased with the number of sessions that separated the fixation maps across all four model layers (V1, V2, V4, IT). A kernel-based maximum mean discrepancy test further confirmed that the empirical distribution of representational distances differed significantly from shuffled controls. These findings suggest that small but systematic shifts in the sampling of a visual scene over time are sufficient to cause representational drift in visual cortex. More generally, these results suggest that subtle changes in behavior over time are inevitable, even in simple tasks, and that these changes may be sufficient to drive representational drift in the absence of intrinsic reconfigurations of neural codes.

**Author summary:** In studies of representational drift in visual cortex, shifts in attention and gaze across repeated stimulus exposures can produce drift-like neural patterns. While the current focus is on contributions of changes in visual sampling to drift, the results raise the possibility that other subtle changes in behavior, many that are hard to quantify, could drive drift in other domains.

## Introduction

Familiar images, such as Vincent van Gogh’s *The Starry Night*, can be recognized across many encounters and viewing conditions. This kind of stable perception of the external world is essential for making effective predictions and planning appropriate actions. For decades, this intuition engendered the assumption that neural responses to the same stimulus would remain highly stable over time. However, longitudinal neural recordings have challenged this view: neural responses to repeated presentations of fixed stimuli can change systematically over days and weeks [1–4]. In a landmark study, Driscoll et al. showed that pyramidal neurons in the posterior parietal cortex gradually shift their preferred spatial locations along a T-maze, even in mice that were already trained to a near-perfect behavioral performance [1]. Decoders trained on neural data from one day therefore generalized progressively worse to data recorded on more distant days. This phenomenon, termed *representational drift*, raises two related questions. First, what mechanisms drive these directional changes that cannot be explained by task changes or random noise? Second, how is the brain able to maintain a stable readout of neural codes in the presence of drift? Recent work has attempted to address these two questions, but beyond identifying drift in multiple brain areas such as parietal cortex [1], olfactory cortex [2], hippocampus [3], and visual cortex [4–7], the answers are still unclear.

Representational drift in visual cortex is especially striking because early visual areas provide a comparatively stable, topographically organized foundation for downstream processing in higher-order cortical areas. However, drift has been observed even in primary visual cortex (V1), despite the need to consistently relay patterns of low-level sensory information from the retina and the thalamus. However, drift in visual cortex has been documented most often in response to complex naturalistic stimuli, such as movies [4, 5] or natural images [6], while simpler tuning properties (e.g., orientation and spatial frequency for gratings) can remain stable over long timescales [4, 5, 7].

A straightforward hypothesis reconciles these findings: complex, naturalistic stimuli afford richer opportunities for active sampling, and this may be accompanied by subtle changes in behavior (e.g., eye movements) that are challenging to fully capture when behavior is summarized at a coarser level. For example, when rodents watch movies, gaze tracking is often omitted from analyses [5] or summarized only as the average pupil position in each experimental session [4]. The challenges associated with eye-tracking arise because, unlike primates, mice lack a fovea and can make non-conjugate eye movements, so accurate mapping of eye position requires specialized calibration and geometric modeling [8]. Similarly, if animals explore different image regions as content becomes familiar, then simply monitoring session-by-session differences in arousal (e.g., pupil size) may also be insufficient. Therefore, unmeasured behavioral restructuring could bias neural measurements toward drift-like patterns. Consistent with this concern, modeling behavioral modulators such as running speed and pupil diameter attenuates drift in several cortical areas [9].

Here, we test whether human gaze during repeated free viewing of naturalistic images changes systematically over experimental sessions, and whether such changes are capable of inducing drift-like shifts in neural representations. We found that fixation-density maps became increasingly dissimilar for sessions separated by greater time intervals. To evaluate the potential impact of these changes on neural codes, each image was gain-modulated using participant-specific fixation profiles and passed through a biologically inspired, weight-fixed deep convolutional neural network (DCNN). Activity patterns in this network exhibited systematic changes over time in all layers, suggesting that altering perceptual sampling is sufficient to induce drift in high-dimensional neural networks that mimic cortical information processing systems.

## Materials and methods

### Longitudinal free-viewing of naturalistic images

Human participants were recruited in compliance with the Institutional Review Board (IRB) at the University of California San Diego. All participants provided informed consent and were compensated 15 USD per hour for participating. Fourteen healthy adults (2 males and 12 females, mean age = 21.2 ± 2.6) with normal or corrected-to-normal vision and intact color vision performed a free-viewing task while being eye-tracked. The study was longitudinal, and participants completed the experiment in 6 separate sessions over an average span of 21± 9.6 days.

No two sessions could be performed on the same day to ensure that gaze-pattern changes were captured over timescales comparable to those used in previous studies of representational drift. Subsequent sessions were separated by 4.2 ± 3.1 days. Each session was divided into 12 separate runs, with each run containing 30 trials. On each trial, a pseudo-randomly selected naturalistic image was displayed on a computer monitor, and the participants were allowed to freely view the image for 5 seconds. The images shown were selected from the DIVerse 2K resolution high quality (DIV2K) dataset [10], because they are more varied and complex than image sets curated for object classification purposes, such as CIFAR-100 [11] or ImageNet [12]. All images were centered and cropped to be of a uniform size (1600 ×1200 pixels, 32.45 ×24.63 degrees visual angle). Different subsets of images were shown across participants. To ensure participants always started by looking at the center of the image, a new trial did not begin until gaze was directed within a circle with a 50-pixel radius around a gray cross-hair that appeared at the center of the screen, which corresponds to approximately 1.05 degrees of visual angle. Once participants fixated on the center of the screen, the gray cross-hair turned black for 1 second signaling the onset of the next image (Fig 1A).

**Fig 1:**
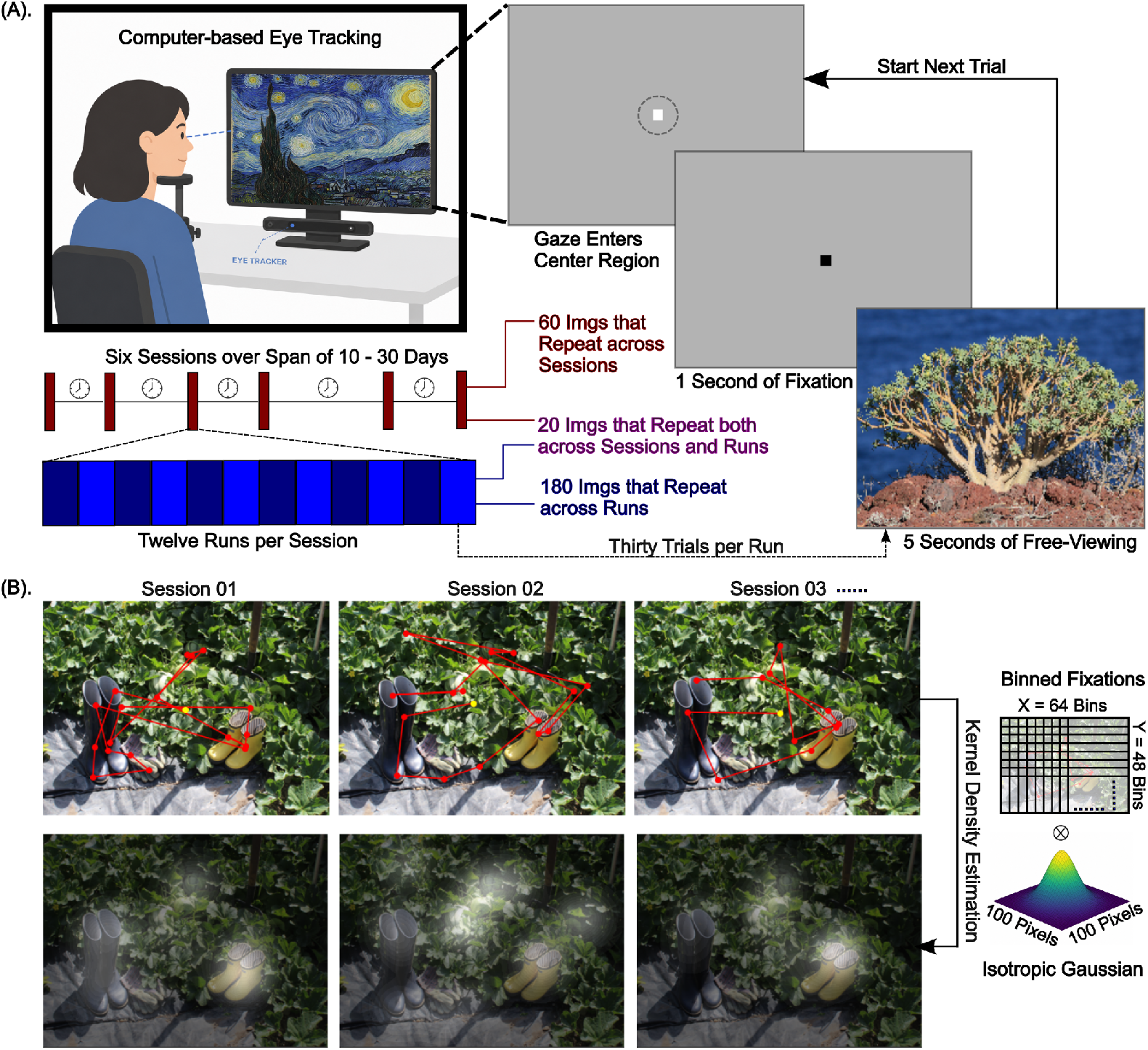
Longitudinal eye tracking. **(A)**: Participants performed a free-viewing task over a span of 6 sessions, where each session contained 12 runs, and each run contained 30 trials. A trial starts when the participant’s gaze entered the central fixation area. The crosshair turned black for 1 second, then a random image appeared for 5 seconds on screen for the participants to freely explore. Images were presented repeatedly at different time intervals, with 60 images repeated once per session, 180 images repeated once every two runs within a given session, and 20 images repeated both across runs and sessions. **(B)**: Gaze trajectories (top) were determined from eye tracking data, and fixation density distributions (bottom) were subsequently derived via kernel density estimation. An isotropic 2D Gaussian kernel with a bandwidth of 100 pixels was convolved against binned fixations and subsequently normalized to derive a 64 × 48 down-sampled fixation density heatmap.

To investigate whether participants systematically changed how they sampled the same visual stimulus over time, images were presented repeatedly throughout the experiment at varying timescales. Out of the 260 unique images that each participant saw during the entire study, 180 images re-appeared once every two blocks for a total of six times within a particular session, 60 of them were shown six times once per individual session, and the remaining 20 images were repeated 6 times both within and across sessions for a total of thirty-six presentations. Participants were not informed about the purpose of the study and were only instructed that they could explore the images freely. At the end of each run, however, they were asked to make an aesthetic rating on 4 additional images that were presented for 10 seconds each. This filler task was created to keep participants engaged throughout the study, and data collected from this aesthetic-judgment task were not used in any subsequent analyses.

### Eye-tracking data

Horizontal and vertical gaze positions were recorded monocularly at 1000 Hz with a desktop mounted EyeLink 1000 Plus [13] as participants freely viewed the naturalistic images. A nine-point calibration was performed at the start of each session, and eye tracking accuracy was validated at the beginning of every run (with recalibration performed as needed).

Gaze data were subdivided into fixations, saccades, and blinks according to EyeLink II version 5.11’s internal detection algorithm, which by default identifies a saccade as an eye movement with a velocity exceeding 30 degrees per second and an acceleration exceeding 8000 degrees per second^2^. Epochs that do not meet these criteria were labeled as fixations. For the purpose of this study, only fixation data were analyzed.

For each participant gaze trajectory was defined as the sequence of fixation locations and durations when viewing a given image. We then generated a fixation density heatmap from the gaze trajectories via kernel density estimation (KDE), using an isotropic Gaussian kernel with a bandwidth of 100 pixels. The KDE was evaluated on a uniform two-dimensional grid spanning the image (64 × 48 bins), yielding a smooth estimate of gaze density that was then normalized to unit area. The resulting fixation density heatmap is thus a discrete probability distribution over binned spatial locations on the given image (Fig 1B). To confirm that results were not specific to the kernel bandwidth selection, we reran all analyses using fixation density heatmaps that were constructed from kernels with smaller (50 pixel) and larger (200 pixel) bandwidths respectively, and the results remained qualitatively unchanged (Fig S3B).

### Quantification of fixation pattern changes

#### Stationary gaze entropy

We first characterized changes in the spatial extent of fixation patterns by calculating the stationary gaze entropy (SGE) [14]. Since entropy is typically highest for uniform distributions, higher values of SGE are associated with more evenly distributed patterns of eye movements and lower values are associated with spatially restricted patterns of eye movements. Each image was divided into two-dimensional grids of size 100 x 100 pixels. SGE is then defined as:

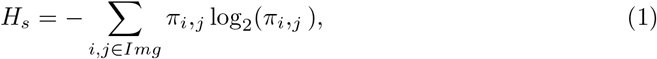

where *π*_*i,j*_ denotes the probability of a fixation falling onto a particular grid *i, j* when viewing that specific image. SGE was calculated separately for each participant and trial, based on the unique viewing profile recorded during each specific image presentation. This metric thus tests whether the spatial extent of fixations changed over time with additional exposure to the same image.

#### Wasserstein distance between fixation density heatmaps

Although SGE captures the dispersion in gaze trajectories, it discards all information about how the patterns changed and is therefore insensitive to changes in fixation patterns if the spatial extent of eye-movements remains constant. To provide a more comprehensive comparison between two sets of gaze trajectories, we calculated the first-order Wasserstein distance (WD), or the Earth Mover’s Distance (EMD), between fixation density heatmaps (Fig 2B). For a participant’s fixation profile when viewing a given image, the EMD between fixation density heatmaps *µ* and *v* collected in sessions *S*1 and *S*2 respectively, is calculated as follows:

**Fig 2:**
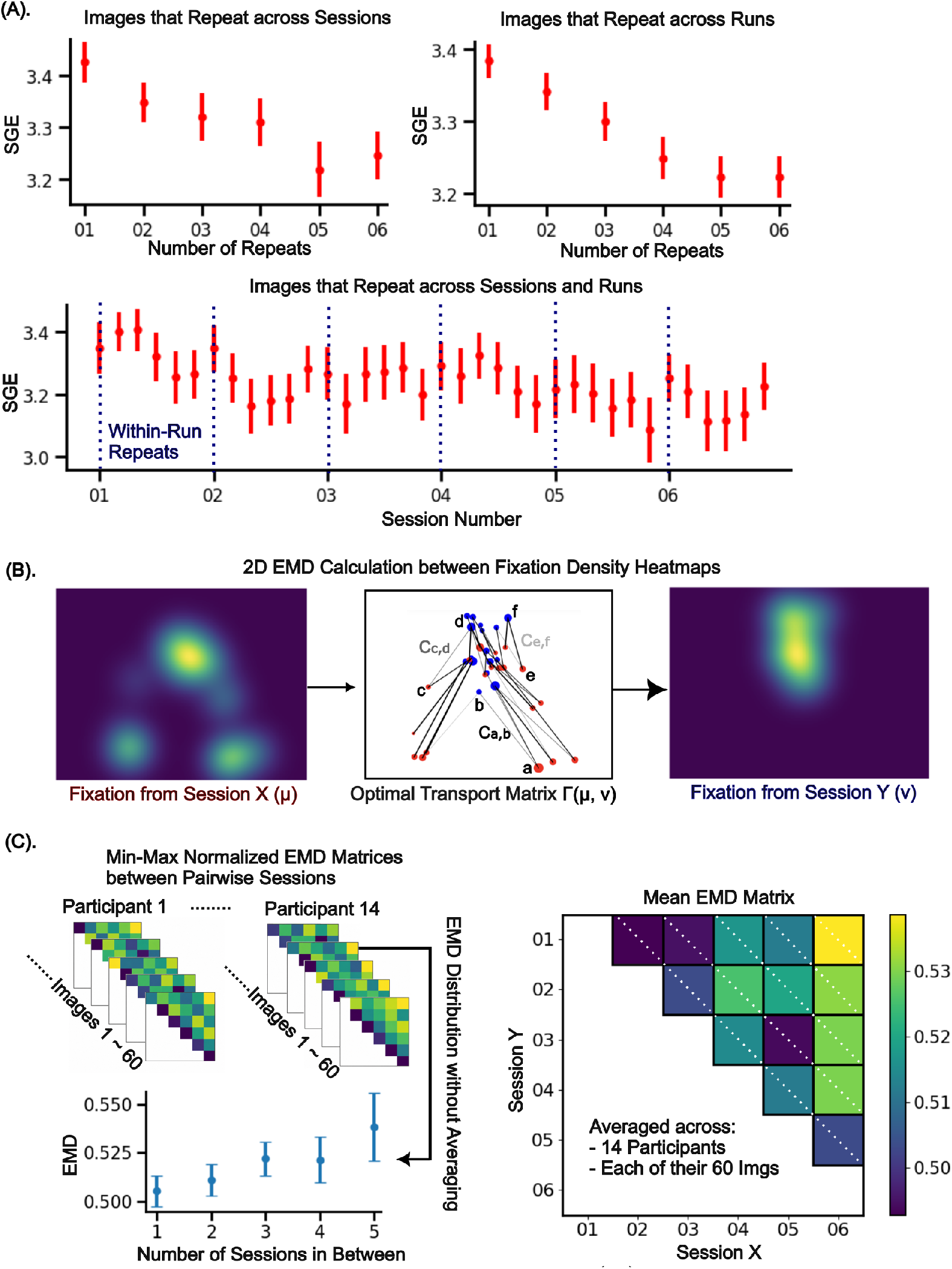
Gaze changes systematically over time. **(A)**: Stationary gaze entropy (SGE) decreased progressively as participants viewed the same image an increasing number of times. The markers represent mean SGE values across all 14 participants and each of the 260 images that they explored. The error bars indicate 95% confidence interval around the mean SGE. **(B)**: Illustration of how the Earthmover’s distance (EMD) is calculated between a pair of 2D fixation density heatmaps denoted as *µ* and *v*. The cost *C*_*i,j*_ of transporting a unit of fixation density from *µ* to *v* is larger (as denoted by darker fonts) if they are separated farther apart in Euclidean space. The amount of mass *γ*_*i,j*_ to be transported between the two locations are determined by the optimal transport matrix, with greater mass denoted by thicker lines in the middle panel. **(C)**: EMDs between fixation density heatmaps were calculated for each session pair, resulting in an EMD matrix for each participant and any given image that they saw (top left panel). The individual EMD matrices were min-max normalized then averaged together to form a single population-level EMD matrix, with the one-off diagonal showing the mean EMDs between fixation density heatmaps that were one session apart, those in the two-off diagonal showing mean EMDs from fixation density heatmaps that were two sessions apart and so on (right panel). Bottom left panel shows the distribution of normalized EMDs between pairs of fixation density heatmaps, grouped by the differing number of sessions in between image presentations. Error bars are 95% confidence intervals around the mean.

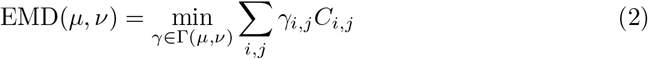

where:

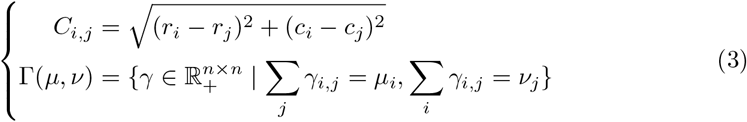

Recall that the fixation density heatmaps are effectively probability distributions on a two-dimensional grid over the original image. *C*_*i,j*_ is thus the cost of moving a single unit of mass from flattened grid index *i* to *j*, and is defined as the Euclidean distance in two-dimensional space between these two grids. This ensures that the spatial properties of fixations are preserved, since a pair of fixation densities that are located farther apart on an image will result in a larger EMD value than those that are closer together in space. The optimal transport plan that captures the flow of fixation density from session *S*1 to *S*2 as determined by the two fixation density heatmaps is denoted as Γ(*µ, v*), where *µ*_*i*_ and *v*_*j*_ are the normalized fixation densities at the grid locations represented by flattened indices *i* and *j* respectively. Therefore, the optimal transport plan ensures that all fixation mass in a given grid from heatmap *µ* is fully accounted for by heatmap *v* after the transportation. The optimal transport plan was obtained and EMD calculations were made using the GeomLoss Library [15], which implements the Sinkhorn algorithm for faster GPU-based computation. A tiny blur parameter (*ϵ* = 10^−10^) was adopted to more closely approximate the exact and unregularized version of EMD calculation.

### Simulation of neural activity in primary visual areas CORnet-S

To investigate whether longitudinal changes in gaze patterns could induce representational drift-like variability in a high-dimensional neural system, we passed images that were masked with the eye-gaze-derived fixation density heatmaps through the biologically-inspired CORnet-S artificial neural network [16]. Note that we did not re-train CORnet-S using our image set and we passed our images through the original pre-trained version of the model. Changes in activation patterns in each layer of the network were then used as a proxy for representational changes in the visual system.

CORnet-S is a convolutional neural network (CNN) with pseudo-recurrence. It consists of four modules that are conceptually analogous to visual areas V1, V2, V4 and IT in the ventral stream, followed by a fully-connected layer for linear classification. Each module is implemented with a canonical bottleneck circuitry that includes sequences of convolutions followed by batch normalizations, channel expansions and compressions, skip connections and a ReLU nonlinearity. Apart from the strictly feedforward V1 module, activity is passed through each of the other three modules more than once using shared weights, where the output of the first pass is added back as input on the next pass, effectively refining the network’s representation at each additional recurrent step. In this implementation, the V2 and IT layers process information twice, while the V4 layer processes information four times (Fig 3A).

**Fig 3:**
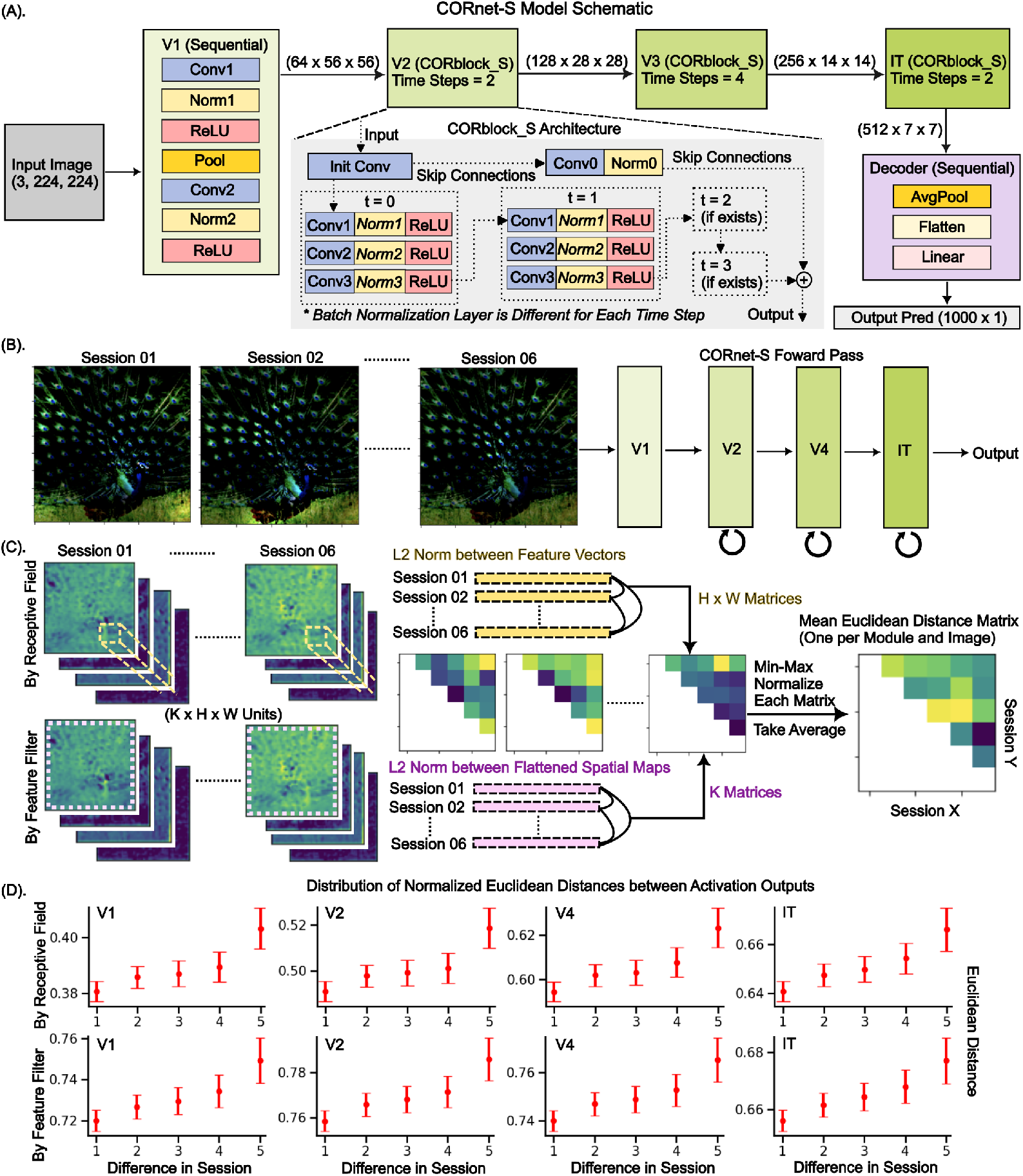
Representational drift-like network activity can be induced by systematically changing fixation patterns. **(A)**: Schematic of CORnet-S architecture, adapted from [16]. **(B)**: For a given image, gain-modulation was applied to the raw image according to the participant’s fixation density heatmaps. This session-dependent modulation creates a set of images that resemble where participants fixated the most across time. These modulated images were then passed through weight-fixed CORnet-S to derive session-specific activation readouts from each module’s output layer. **(C)**: For any given module, response vectors from its activation readouts can be defined as either a feature vector across many receptive field locations (top row, yellow) or a flattened spatial activation map across different feature selective kernels (bottom row, pink). The Euclidean distances between response vectors derived from different session pairs populate a set of session-by-session Euclidean distance matrices. The matrices were then min-max normalized and averaged to form a single mean distance matrix for each image. This represents the average network-level perturbations caused by differing sampling of the input image. **(D)**: The distance between network activations increased as input images were modulated by empirical fixations that were further separated by sessions. The distribution includes distances derived from all images that were repeated across sessions, and error bars represent 95% confidence interval around the mean.

Although there are many candidate networks, we intentionally selected CORnet-S for its interpretability and functional similarity to the brain. Despite its comparatively shallow architecture, the pseudo-recurrent structure in CORnet-S allows it to perform on par with some of the more classic deep models such as ResNet-50 and VGG19. More importantly, CORnet-S was the leading model when evaluated against Brain-Score [17], a composite metric that measures how closely a model’s activity resembles that of the primate brain when the animals and models perform similar tasks. Since we aimed to investigate how behaviorally-induced changes can affect neural readouts, CORnet-S strikes an ideal balance between machine learning performance and biological validity.

#### Fixation-density-modulated images as model input

To assess how shifts in gaze patterns over time impacted neural representations, the input images passed into CORnet-S were masked with their associated fixation density heatmaps from one of the sessions in the empirical study. The mask *M*_*norm*_ was defined as:

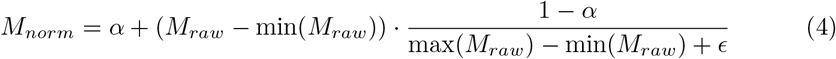

For a given session where a participant explored a specific image, *M*_*raw*_ refers to the original fixation density heatmap derived from that participant’s eye-tracking data. Min-max normalization was applied to *M*_*raw*_ with a lower bound parameter *α* that determines the baseline visibility after masking. When *α* = 0, unvisited locations on the image were zeroed out (i.e. black). Setting *α* to a small or moderate value therefore allows the background to remain slightly visible, which is more biologically realistic and is beneficial for CORnet-S as it provides some global context of the image outside attended regions. Our main analyses were run with *α* = 0.2, but to evaluate the stability of the results, we quantified model outputs under additional values of *α* ∈ {0.0, 0.4, 0.8} and the results remained qualitatively unchanged (Fig S3C).

The original image *I*_*original*_ was then linearly scaled by the mask *M*_*norm*_ as

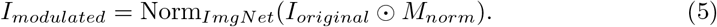

To ensure that the modulated images were compatible with the CORnet-S architecture and consistent with the ImageNet image statistics that were used to train CORnet-S, the modulated image was spatially transformed to be of size 224 × 224 pixels, followed by normalization to have channel-wise mean *m* = [0.485, 0.456, 0.406] and standard deviation *σ* = [0.229, 0.224, 0.225]. The final product is a fixation-modulated, size-matched, and normalized image *I*_*modulated*_ that was compatible with the CORnet-S model (Fig 3B), with the different *I*_*modulated*_ images reflecting the varying set of regions that a participant sampled in each session.

#### Quantification of model output

With the model input reflecting how a participant differentially scanned a given image over time, the final output of each module in CORnet-S is thus a proxy for neural responses in each visual area. For any given input image, the activation output of each module *p* has shape [*H*_*p*_ ×*W*_*p*_ × *K*_*p*_], where the exact dimensions are determined by the CORnet-S architecture. Therefore, each unit in that module represents the activation of a specific feature filter (*k*) that is spatially localized to a particular part (*h, w*) of the image after convolution. To quantify session-by-session changes in network activity, a pair-wise distance matrix *D* was calculated, with each of its elements *D*_*i,j*_ as the Euclidean distance between two response vectors 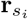 and 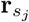:

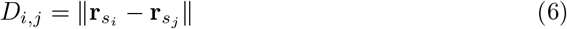

where *s*_*i*_ and *s*_*j*_ are session numbers associated with the empirical eye-tracking data that were used to modulate input images to the model.

The response vector **r**_*s*_ can be defined in two different ways: either as a feature vector of shape [*K*_*p*_ × 1] for a particular spatial location (*h, w*), or as a flattened spatial activation map of shape [(*H*_*p*_ ×*W*_*p*_) ×1] for a select feature filter *k*. The former allowed investigation of activity based on each unit’s spatial receptive field, while the latter allowed investigation of activity based on each unit’s feature selectivity profile. For the purpose of our study, we were interested in the population-level stability of network representations, and thus did not analyze the distance matrices individually. We first min-max normalized each pair-wise distance matrix *D* derived from different sets of response vectors **r**_*s*_. We then averaged across all distance matrices to derive a single network-wide pair-wise distance matrix (Fig 3C). Depending on how the individual response vectors were defined, we ended up with either a single matrix *D*_*RF*_ (pooled across all spatial receptive fields) or *D*_*FF*_ (pooled across all feature selectivities) for each image and participant.

#### Nonparametric test of distributional changes

To determine if pair-wise distance matrices *D*_*RF*_ and *D*_*FF*_ contained meaningful internal structure that was distinct from random noise, we used the maximum mean discrepancy (MMD) as a nonparametric two-sample test statistic for quantifying differences between distributions [18]. The empirically derived distance matrices served as one set of distributions, while those derived from resampled gaze data (see **Resampling as null distribution**) served as the other. Thus, the MMD statistic characterizes the distance in mean embeddings between these two distributions when projected into a reproducing kernel Hilbert space, with larger values indicating greater dissimilarity between the two datasets (Fig S4A). For details see Supporting Information: Maximum mean discrepancy test details.

### Resampling as null distribution

In addition to parametric statistical tests, we also used resampling to assess the significance of the results of all analyses described above (i.e., SGE, EMD and CORnet-S activation patterns). For each participant and image, we pooled their gaze trajectories by concatenating both the location and the duration of individual fixations across all six sessions. We then shuffled these fixations and resampled from the pool, resulting in a new set of fixation profiles across repeated exposures to a given image (Fig S1A top). This resampling method ensured that the simulated eye-tracking data were ecologically valid, as each individual segment of fixation was directly derived from the empirically recorded gaze positions. In addition, the shuffling component disrupted any longitudinal trends of shifting fixation patterns, thus providing a baseline to evaluate against our empirical results. We compared the summary statistics (such as slope of SGE over sessions) derived from the empirical gaze data to the distribution of those from the 500 resampled datasets. The p-value from this non-parametric test determines whether the empirical results are distinguishable from the resampled null.

To determine if the constraints of this resampling method introduced any biases, we also ran the same non-parametric tests on session-shuffled gaze datasets as a more conservative null distribution. For details see Supplemental Information: Resampling method caveats and validation.

## Results

### Stationary gaze entropy decreases as a function of repeated image presentation

With repeated presentations of an image across sessions, SGE significantly decreased (ordinary least squares linear regression, slope=-0.037, p*<*0.05). Similar effects were present for images that repeated on a shorter time scale within a single session (slope=-0.034, p*<*0.05) and for images that repeated both within and across sessions (slope=-0.005, p*<*0.05) (Fig 2A). To ensure that the decrease in SGE was not caused by participants spending less time fixating on an image towards the end of the experiment due to looking off-screen or blinking more frequently, we ran the same regression analysis but only retained trials where participants spent more than 80% of their time fixating on the image, which accounted for 80.4% of all across-session observations. After removing trials with lower levels of fixation, the decrease in SGE across sessions remained statistically significant (slope=-0.047, p*<*0.05, Fig S3A). A non-parametric permutation test against the distribution of slopes derived from our resampled gaze datasets further confirmed that this across-session decrease in SGE in the full dataset was not due to random chance (resampling-based non parametric test, two-tailed p*<*0.002, Fig S1B).

Although there was variability across images and participants, the population-level regression results imply that, on average, the gaze distribution became more concentrated as participants became more familiar with image content. This is consistent with previous studies that found reduced active visual exploration in participants with repeated viewing of the same naturalistic scenes [19]. However, prior experiments were almost exclusively task-driven, and participants were asked to memorize or visually search for specific objects [19, 20]. Under a task-based context, participants are more likely to consistently scan the most informative parts of the image to optimize task performance. Our results demonstrate that exploration is also attenuated even in the absence of clearly defined goals to guide behavior, and that gaze distributions naturally become more localized over time.

### Gaze distribution shifts systematically over time

To better capture changes in the spatial pattern of eye-movements, we computed the EMD between fixation density heatmaps across pair-wise sessions for each image and participant. To compare across observations, we performed min-max normalization on the computed EMDs within each image before averaging across images and performing a population-level regression analysis. This approach ensured that the EMD values were always scaled between 0 and 1 according to the overall distribution of EMDs within any particular image, preventing images with high EMD values from dominating the results. On average, the EMD between fixation densities that were separated further apart in time were larger than those that were closer together, as indicated by a systematic increase in EMD with increasing separation between sessions (ordinary least squares linear regression, slope=0.007, p*<*0.05) (Fig 2C).

Like SGE, the EMD is sensitive to gaze dispersion, but it also captures changes in the spatial patterns of fixations. Thus, even though changes in SGE and EMD across sessions should generally be correlated, they can diverge if only certain aspects of gaze changed over time. For example, if fixations are uniformly dispersed over a fixed area but the centroid of the fixations gradually shifts in the same direction over time, then SGE will stay the same but EMD will steadily increase. Our observation that both SGE and EMD change as a function of separation between sessions is consistent with directional drift in both the spatial extent of fixations and the center of mass of those fixations as participants explore increasingly familiar images.

While our regression analysis revealed a significant increase in EMD across sessions separated further in time, the variance explained is small (R2=0.0012) due to high variability between observations, potentially because each participant viewed a different subset of images with varying low-level statistics and semantic complexity. Therefore, we also ran a non-parametric test by performing the same regression analysis but on resampled null datasets where longitudinal associations between gaze were disrupted (see **Resampling as null distribution**). We found that of the 500 resampled datasets, the slope of EMD across separation between sessions was consistently smaller than that of the slope observed in the intact data, and the null distribution was centered around zero as expected (resampling-based non parametric test, two-tailed p=0.004, Fig S1C). Together, these results confirm robust, systematic changes in the gaze patterns over time.

### CORnet-S activations show representational drift-like trends

To test whether systematic drift in the sampling of visual information is sufficient to induce drift in neural signals, we passed the images through CORnet-S while masking them using fixation densities derived from the empirical data. Thus, the model received a modulated visual input where more frequently visited features were emphasized and less frequently visited were attenuated. We again performed a linear regression on the Euclidean distances between pairs of network activations to test whether they deviated more from one another as the input images were modulated by gaze data collected in further separated experimental sessions. We found that across all network modules, activations became more dissimilar over time as the network processed images modulated by the empirical gaze data (Fig 3D; ordinary least squares linear regression, slope=0.0041 for module V1, p*<*0.05; 0.0049 for module V2, p*<*0.05; 0.0055 for module V4, p*<*0.05; and 0.0053 for module IT, p*<*0.05).

We followed up with non-parametric resampling tests, where the same analysis was run using images modulated by masks generated from the resampled null dataset. In contrast to the empirical data, network activations did not grow systematically more dissimilar as a function of difference between sessions when using randomized datasets, and the slope obtained from the regression analysis was smaller than the slope associated with the empirical data in all four network modules (resampling-based non parametric test, two-tailed *p <*0.002 for all layers, Fig S2). Furthermore, a 1000-fold permutation test on the MMD statistic of the flattened Euclidean distance matrices revealed a statistically significant difference in the representational geometry of network activations induced by empirical versus resampled gaze-modulated inputs (p*<*0.05 for all layers when grouped by receptive fields as in Fig S4B; p*<*0.05 for V2 and V4, p=0.06 in V1 and p= 0.13 in IT when grouped by feature filters as in S4C).

The comparisons between empirical and randomized data demonstrate that systematic changes in gaze behavior can induce directional drift in the activity patterns of a biologically inspired neural network.

## Discussion

Representational drift refers to systematic changes in neural responses to the same stimulus over days and weeks. Our results show that human gaze patterns do not remain stable across repeated free viewing of the same stimulus. Instead, SGE decreased and the EMD between fixation-density maps increased with greater separation between sessions. While SGE strictly measures the spread of gaze allocation, EMD also accounts for the spatial distribution of gaze. The observation that both metrics change over time suggests that fixations became increasingly concentrated and that participants gradually shifted their gaze toward different parts of the image. Note that this can be achieved in at least two ways: by exploring a similar set of locations on an image but with higher precision, or by reducing unnecessary exploration and only fixating on a few highly salient features. Together, these patterns indicate that this behavioral drift changes the sensory input received by the visual system over time. Accordingly, when images were masked using empirically-based fixation-density heatmaps, weight-fixed CORnet-S activations became increasingly dissimilar across sessions in all four modules (V1, V2, V4, IT). Because each module maps onto corresponding stages of the primate ventral stream, systematic changes in input sampling may propagate through hierarchical visual processing and remain detectable in later, more abstract layers.

A central assumption in many drift studies is that behavior is stable once animals are over-trained, so neural changes reflect internal representational restructuring. This interpretation often assumes that sensory input remains constant across sessions. However, even when task accuracy is stable, behavior can continue to adapt in ways that do not alter measured performance. In rodent navigation, for example, coarse body-position tracking can appear stable while finer kinematic variables (head tilt, whisking frequency) still change [21, 22]. If such behavioral changes are systematic, corresponding changes in sensory input could propagate through neural systems and manifest as drift-like neural dynamics, even in areas that are considered distal from early sensory areas. It is also notable that in the present task, systematic gaze restructuring emerged even in passive free viewing. Task-driven viewing would likely amplify this effect, as participants may shift from broad exploration towards efficient scanning of just the most task-relevant features. After training, gaze may therefore continue to reorganize even when measured accuracy is stable, providing another source that could drive neural drift in sensory processing tasks.

When CORnet-S readouts were summarized via feature vectors pooled across space, baseline dissimilarity between conditions increased in later layers, while the session-dependent increments in dissimilarity were relatively stable across layers. This pattern likely reflects nonlinear accumulation of input perturbations in a feedforward neural network, as local gaze-induced changes in image statistics are amplified when activity passes through successive modules. This does not necessarily contradict findings that later layers can be more invariant for classification under some transformations [23], because representational geometry and classification invariance are distinct properties. For example, prior fMRI work with eye tracking reported stronger gaze-related representational differences in early visual cortex than in IT [24]. This divergence from our simulation results may also reflect the absence of long-range feedback and top-down attention mechanisms in CORnet-S, which stabilize representations in biological vision. However, when readouts were instead defined by spatial locations pooled across feature filters, this layer-wise amplification of baseline dissimilarity was weaker, suggesting that gaze modulation affected feature-space organization more than coarse spatial topology.

A limitation of the current empirical study is that each participant viewed a different image set, restricting our ability to test how image statistics such as low-level salience or top-down interest might shape systematic changes in sampling. Image complexity likely influences both the magnitude and consistency of gaze shifts: sparser scenes may support fewer stable fixation strategies, whereas denser scenes might encourage continued refinement of sampling. Future work should use a shared image battery with controlled variation in complexity to better determine if common factors drive drift across individuals.

Our modeling approach also has limitations. CORnet-S provides a useful but indirect proxy for biological responses, and convergent evidence will require simultaneous neural recording and eye tracking in an animal model system that has more human-like gaze patterns. We also held model weights fixed, which matches the absence of explicit task goals that our participants experienced. As a result, we are unable to disentangle implicit from explicit causes of changes in gaze, which might be learned passively via repeated exposure or actively due to changes in the perceived relevance of different parts of an image. However, because drift-like effects were already present under this conservative weight-fixed model, allowing weight updates would likely further increase representational change. Finally, masking images with fixation-density maps preserves where participants looked most often but discards within-trial temporal structure. Follow-up work could use recurrent dynamics in CORnet-S to examine how network activity unfolds over the course of each trial with gaze trajectories as input.

The present findings do not argue against the existence of intrinsic representational drift driven by mechanisms such as synaptic turnover [26], learning-related plasticity [25, 27, 28], changing internal states [9], and efficient coding [29]. Instead, they offer a complementary account: systematic changes in how sensory inputs are sampled can produce drift-like neural patterns without requiring the reorganization of internal circuitry. Contributions from changes in perceptual sampling may thus help explain why drift is more readily observed with complex naturalistic stimuli, as such stimuli provide richer context that encourages more exploration compared to simple stimuli such as oriented gratings. Although our experiment only tested this hypothesis within the domain of visual processing, the same argument should apply to other sensory systems, including multi-sensory and memory-related systems such as the hippocampus. Our results therefore suggest future studies should employ detailed behavioral monitoring as a prerequisite, with careful consideration of even subtle changes in behavior that are challenging to quantify. When such behavioral stability cannot be established, regressing out behavioral modulators and testing whether drift is attenuated is a critical control, particularly in tasks involving complex naturalistic stimuli where many strategies might support equivalent accuracy.

## Conclusion

Our study showed that active sampling of the external environment can change systematically over time even in the absence of a goal-directed task. Such behavioral changes can induce perturbations in sensory inputs that increasingly deviate over time and are amplified as they propagate through a processing hierarchy. The net result is population-level activity changes that resemble directional representational drift but is purely driven by changes in patterns of input. Future studies will need to take into account potential longitudinal changes in behavior when investigating the properties of neural code re-structuring in the context of studying representational drift. Most importantly, our results further suggest that drift is a normal property of neural codes, as non-stationary sampling of the environment occurs in natural vision and these changes will inevitably impact the long-term stability of neural codes.

## Acknowledgements

This work was supported by the National Eye Institute under grant number **R01EY025872**.

## Supporting information

### Maximum mean discrepancy test details

To calculate the MMD statistic between a set of empirical and a set of resampled distance matrices, a kernel matrix *K* that captures the similarity between every pair of data points in these two distributions is required. Each distance matrix from the two datasets was flattened into a single vector, and the Euclidean distance between any pair of such vectors was projected into high-dimensional similarity space via a Radial Basis Function (RBF) kernel. For example, given two flattened distance matrices **d**_*i*_ and **d**_*j*_, their similarity in kernel space is calculated as:

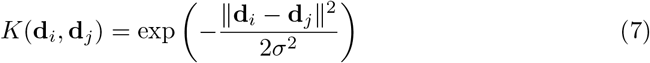

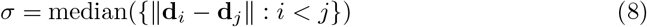

where the kernel bandwidth *σ* was selected to be the median Euclidean distance between all pairs of flattened distance matrices. This ensures that the scale of the kernel matrix matches that of the original data distribution, supporting optimal sensitivity when discriminating between the two samples.

After populating the kernel matrix *K* with all combinations of **d**_*i*_ and **d**_*j*_ from the two datasets, the unbiased MMD statistic was calculated as:

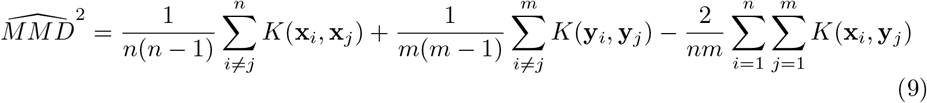

where 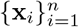 are the set of flattened distance matrices from the empirical dataset, and 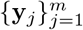 are those from the resampled dataset (Fig S4A). To test the significance of this MMD statistic, group labels indicating whether distance matrices belonged to empirical or resampled distributions were shuffled across the combined dataset to generate 1000 permutation pairs, and the corresponding p-value was calculated based on how the original MMD statistic compares to that of these shuffled pairs containing mixed samples. The resulting p-value therefore indicates whether the distribution of empirical distance matrices is significantly distinguishable from that of the resampled.

### Resampling method caveats and validation

One limitation of our primary resampling method is that we cannot preserve both the number of fixation segments and the total fixation duration to be strictly the same as the empirical data. For example, if a participant had only a single 4-second long fixation when viewing an image in one session, this fixation segment might be re-assigned to another session and concatenated with other segments to create a fixation trajectory that exceeds the 5-second trial duration. Theoretically, this doesn’t affect downstream analysis, since the calculations for SGE and EMD are based on spatial distributional properties of gaze rather than the absolute timing. However, to confirm that this caveat did not bias our statistical tests, we ran a simpler non-parametric control test by keeping the gaze content within each session intact while shuffling the session labels of each image 500 times (Fig S1A bottom). We then evaluated our empirical summary metrics against a null distribution that was derived from session-label-shuffled datasets free of fixation re-distribution and concatenations. The results remained statistically significant, confirming that the resampling artifact did not bias our primary findings (Fig S1B, S1C and Fig S2).

**Supplementary Figure S1.**
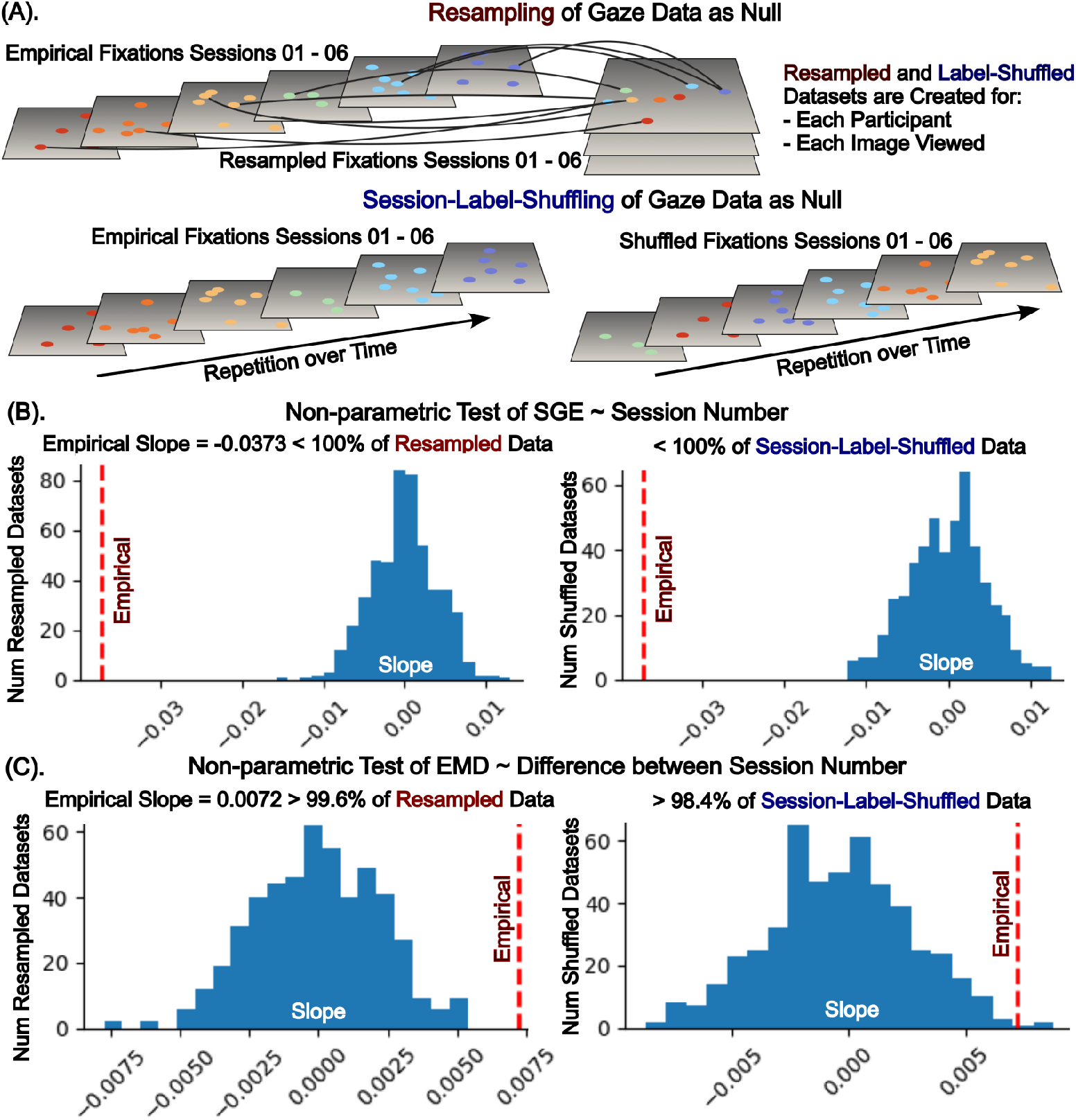
Null gaze datasets do not demonstrate systematic changes in fixation profiles over time. **(A)**: Top - chunks of fixations were randomly selected and re-distributed across different sessions to create a resampled gaze dataset. Bottom - fixation profiles were left intact while the session labels were shuffled. **(B)**: Across all resampled (left) and session-label-shuffled (right) datasets, SGE remain stable as the number of times an image was presented increased, with slopes centered around zero and consistently smaller than that of the empirical dataset. **(C)**: Across the vast majority of resampled (99.6%, left) and session-label-shuffled (98.4% right) datasets, EMD between fixation density heatmaps remain stable as the number of sessions between image presentations increased, with slopes centered around zero and smaller than that of the empirical dataset.

**Supplementary Figure S2.**
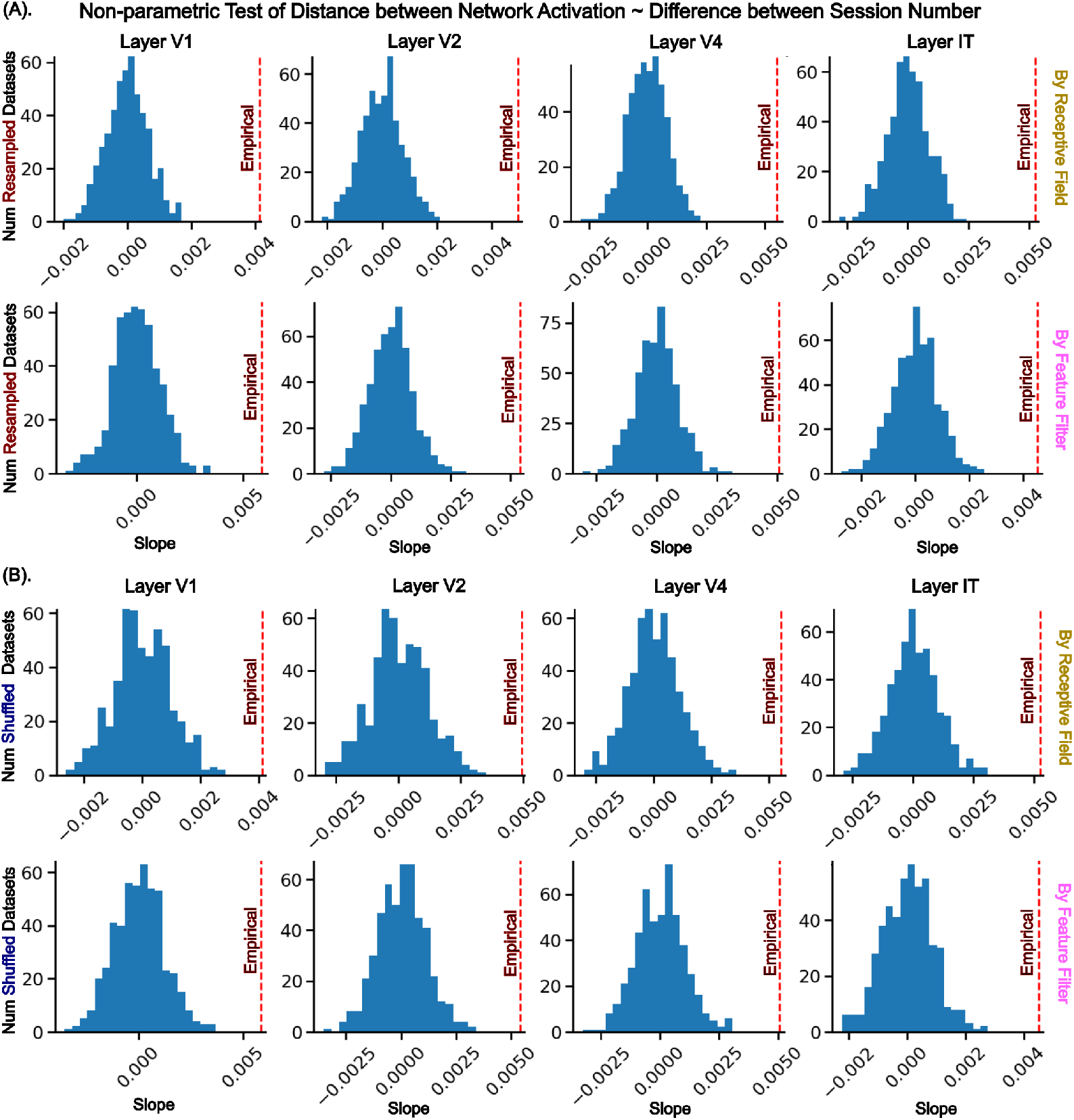
Null gaze datasets do not induce representational drift-like neural patterns when passed through CORnet-S. **(A)**: Across all resampled datasets, the dissimilarity between CORnet-S network activations remain stable as the pairs of input images were modulated with gaze data that were sampled from further separated sessions, as indicated by slopes centered around zero and consistently smaller than that of the empirical. **(B)**: The same was true for all session-label-shuffled datasets.

**Supplementary Figure S3.**
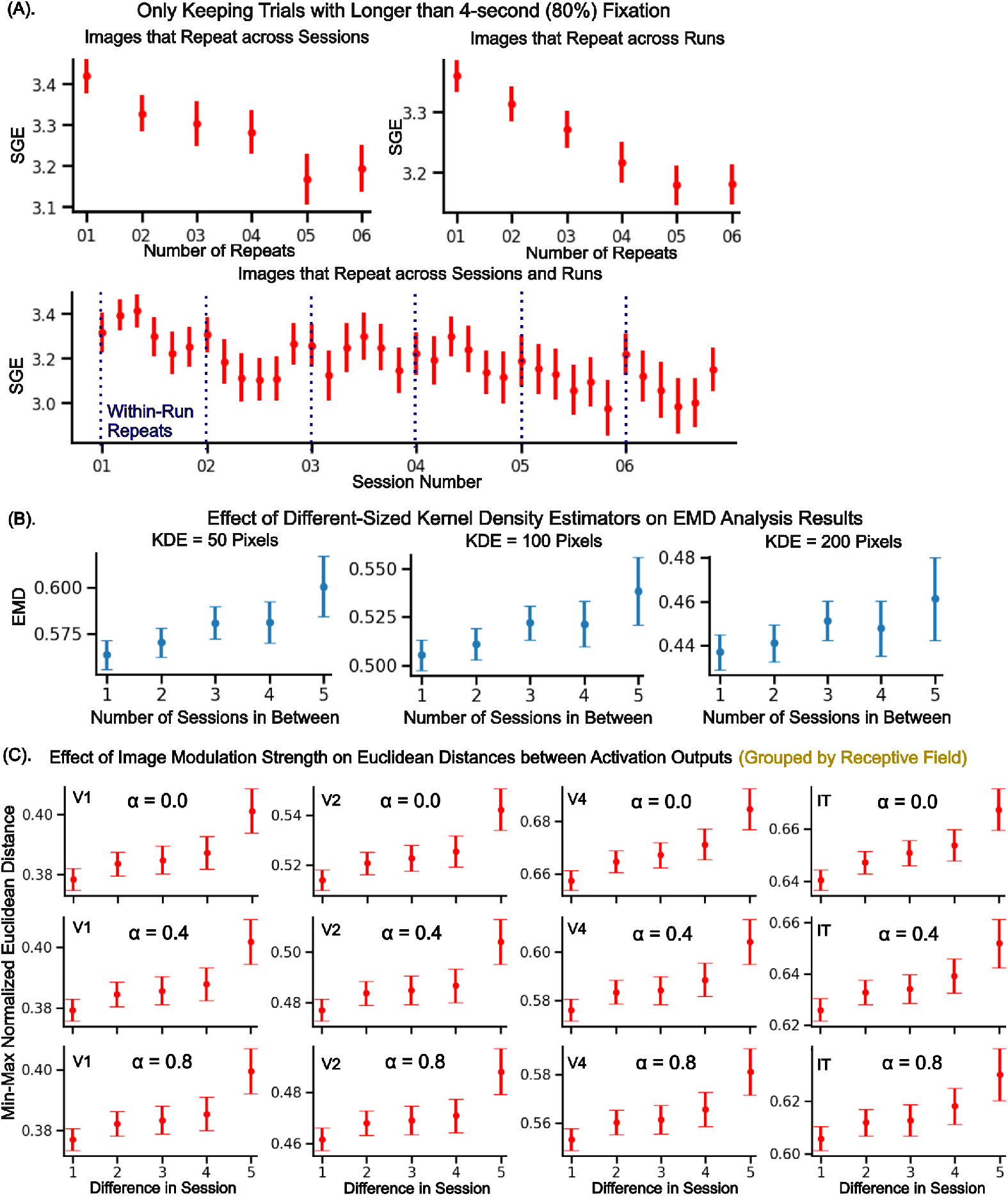
The choice of parameters in the processing and analysis pipeline does not qualitatively alter the primary results. **(A)**: SGE as a function of image repeats, excluding trials where the amount of time spent fixating was less than 80%. The markers represent mean SGE values across all 14 participants and each of the 260 images that they explored. The error bars indicate 95% confidence interval around the mean SGE. **(B)**: EMD as a function of number of sessions between image presentations, where fixation density heatmaps were created under different KDE sizes. **(C)**: Select examples of dissimilarity between CORnet-S network activations as a function of number of sessions between sampled gaze data, where the input images were modulated at different strengths according to various values of *α*.

**Supplementary Figure S4.**
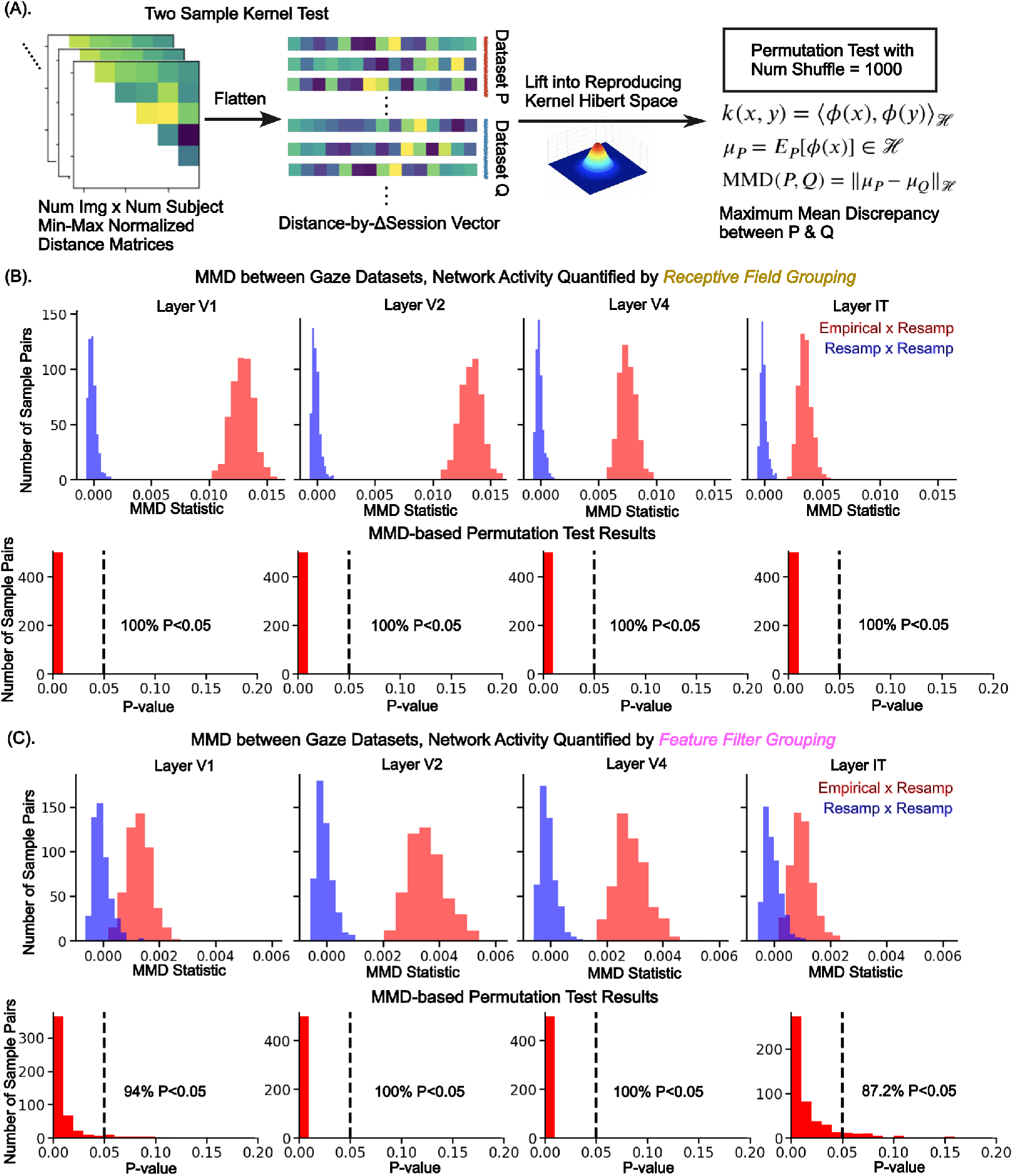
MMD-based two-sample permutation test. **(A)**: Schematic of the testing procedure. **(B)**: For network activation distance matrices quantified via grouping units by their receptive fields, the MMD between those derived from the empirical and the resampled null datasets (top red), as well as the permutation test results with 1000-fold shuffling (bottom). **(C)**: Same as **B** but for network activation distance matrices quantified via grouping units by their feature selectivity profiles.

## Notes

### Competing Interest Statement

The authors have declared no competing interest.

### Summary of Updates

Updated co-senior author order, moved Mikio Aoi to last author.

